# Towards Spectral Variation Analysis: A Data Quality Framework for Non-Targeted Methods

**DOI:** 10.1101/2024.11.27.625758

**Authors:** Kapil Nichani, Steffen Uhlig, Victor San Martin, Bertrand Colson, Karina Hettwer, Ulrike Steinacker, Heike Kaspar, Petra Gowik, Sabine Kemmlein

## Abstract

Non-targeted methods (NTM) require robust methods for comparing spectral data for reliable classification and identification. Traditional approaches using match factors reduce complex spectral relationships to single values, limiting their utility in quality assurance. This study presents an evaluation of spectral comparison methodologies, contrasting classical Mahalanobis distance (MD) with neural network approaches, namely, neural classification distance (NCD). Using matrix assisted laser desorption ionization-time of flight (MALDI-TOF) mass spectrometry data from bacterial isolates, we systematically assessed these methods across varying levels of spectral variability. The MD approach exhibited consistent performance under controlled conditions but showed limitations with increasing spectral complexity. In contrast, the NCD demonstrated adaptability across all scenarios, revealing its capability in handling complex spectral relationships. Through this exemplary example, we present the mathematical framework for quantifying spectral variations and establish criteria for method selection in different analytical scenarios. This work provides a foundation for proposing data quality metrics in NTMs and offers practical implementations for routine quality assurance. The methodology developed here extends beyond mass spectrometry applications and contributes to the broader field of analytical quality control in complex spectral analysis.

## 1 Introduction

Non-targeted methods (NTM) have emerged as a pivotal analytical approach in environmental monitoring [1,2], food safety [3,4], and forensic applications [5,6]. The methodology relies on comprehensive fingerprinting, particularly using measurement data generated by next generation sequencing (NGS), spectroscopy or mass spectrometry, to characterize unknown samples without a priori knowledge of their composition [7]. Sticking to the case of spectral data hereafter, the statistical classification of these spectral patterns into predefined categories (e.g., A or B) can be represented mathematically as:

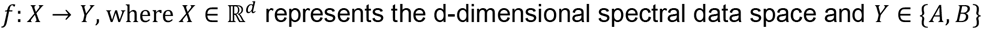

The reliability of such classifications fundamentally depends on spectral data integrity. Poor quality data inevitably leads to misclassification, with potentially serious consequences in critical applications such as food safety [8] and forensics [9]. Both poor quality training data and “out-of-population” data during inference can equally undermine model performance through subtle degradation mechanisms that often escape standard quality control measures. Despite this criticality, quantitative methods for comparing spectral data remain underdeveloped [10–12]. Specifically, the question regarding how to examine variations in spectra arising from (a) time or instrumental factors within single-sample measurements, under repeatability conditions and (b) inter-laboratory differences across multiple samples, under reproducibility conditions. This study attempts to address this gap by proposing new methodologies for spectral data comparison and developing robust metrics for quantifying these spectral variations that will provide a foundation for implementing comprehensive quality assurance protocols in NTM applications.

## 2 Problem statement

In traditional analytical measurements, measurement error *ε*_*i*_ for a single measurement point *x*_*i*_ is typically characterized as a univariate value *ε*_*i*_ = *x*_*i*_ − *μ*_*i*_ and 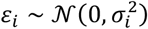 (or some parametric or non-parametric distribution) [13]. Measurement error theory further allows hierarchically “decomposition” of the error components, e.g., 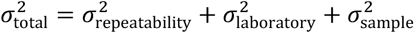.

However, spectral data presents a more complex scenario, where we have a multivariate measurement. Let us define a spectrum ***S*** as an ordered set of intensity measurements across d dimensions: ***S*** = {*x*_1_, *x*_2_, … , *x*_*a*_} where *x*_*i*_ ∈ ℝ^*d*^. If ***μ*** is a true spectrum, then analogously, ***ε*** = ***S*** − ***μ*** and ***ε*** ∼ 𝒩(**0**, Σ), where Σ is the covariance matrix.

The challenge with spectra is already becoming evident here as acquiring a “true” spectrum may be challenging or virtually unattainable. Just like in univariate case, spectral measurement error also exhibits a hierarchical complexity depending on the experimental conditions. At the most fundamental level, variations arise even when spectra are acquired from the same sample, using identical instrumentation, within a single measurement session - representing the inherent instrumental repeatability. The error magnitude typically increases when considering reproducibility: measurements of the same sample across different times, capturing instrument drift and environmental fluctuations. Further complexity emerges in inter-laboratory comparisons, where systematic differences in instrumentation, calibration procedures, and operator practices contribute additional variance components. The highest level of variability manifests when comparing measurements across different sample preparations or batches, incorporating sample heterogeneity and handling effects into the error structure.

The fundamental problem can be formulated as finding a function φ that maps two spectra *S*^1^ and *S*^2^ to a set of characteristics C that describe their relationship: *φ*: (***S***^1^, ***S***^2^) **→ C**.

The challenge lies in determining φ such that it satisfies: **C** = *φ*(***S***^1^, ***S***^2^) where **C** ∈ ℝ^*d*^. We seek a generalized concept for measurement error for NTMs and this work is an attempt to propose in this direction.

## 3 Distances for spectral comparison

The hierarchical variations in spectra directly impact method performance characteristics (α), which can be expressed as a function of Q representing data quality metrics, N is the sample size, and σ denoting the variation in the measurement, *α* = *g*(*N, Q, σ*). Both Q and σ can be quantified through “distances” - mathematical measures that determine similarity or dissimilarity between spectra.

To determine appropriate distance measures for spectral comparison, we must consider both the spectral characteristics (peak positions, shapes, and relative intensities) and the underlying physical phenomena causing variations. In the following, we propose two complementary approaches.

### 3.1 Mahalanobis distance (MD)

We examined spectral differences through classical multivariate statistical lens using the Mahalanobis distance (MD). Consider a set of n reference spectra represented by the data matrix ***X*** ∈ ℝ^*n*×*q*^, where each row represents a spectrum ***X***_***i***_ ∈ ℝ^*q*^, *i* = 1, … , *n*. For mass spectrometry data, *q* represents the number of mass-to-charge (m/z) channels, and each element *x*_*ij*_ represents the intensity at the jth m/z value in spectrum i. The Mahalanobis distance *MD*(***X***_***i***_) between a spectrum ***X***_***i***_ and the distribution of reference spectra is defined as:

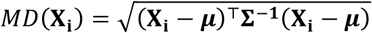

where, ***μ*** ∈ ℝ^*q*^ is the mean spectrum of the reference set and **Σ** ∈ ℝ^*q*×*q*^ is the covariance matrix, which can be calculated as:

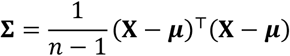

The covariance matrix describes the correlations between multiple variables. It is a square matrix with dimensions equal to the number of variables, and its entries represent the covariances between the variables. The diagonal entries of the covariance matrix represent the variances of the individual variables, which measure the dispersion or spread of the data around the mean. The off-diagonal entries represent the covariances between pairs of variables, which measure the extent to which the variables are linearly related. The expected value of the Mahalanobis distance *E*(*MD*) for *d* degrees of freedom can be derived as 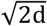:

### 3.2 Neural classification distance (NCD)

Alongside MD, in this study, we introduce the Neural Classification Distance (NCD), which quantifies spectral variations through neural network-based classification probability. Let *f*_*θ*_: ℝ^*q*^ **→** [0,1] be a neural network with parameters *θ* that maps input spectra to classification probabilities (*p*). This network function can be decomposed as: *f*_*θ*_ = *g* ∘ *ϕ*, where *ϕ*: ℝ^*q*^ **→** ℝ^*k*^ is the internal transformation through hidden layers and *g*: ℝ^*k*^ **→** [0,1] is the final sigmoid activation function. The internal transformation *ϕ* implements a non-linear mapping through M hidden layers:

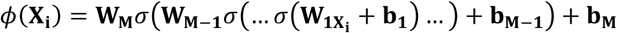

Where, **W**_**m**_ are the weight matrices, **b**_**m**_ are bias vectors and *σ*(⋅) is a non-linear activation function. The NCD is then defined as:

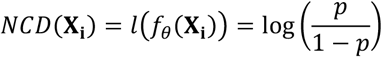

Where *p* = *f*_*θ*_(***X***_***i***_) is the network’s output probability for spectrum i and *l*(⋅) is the logit transformation.

Neural network classifiers would be trained to distinguish between spectral classes, incorporating both repeatability and reproducibility aspects. These logit-transformed probabilities serve as a natural distance metric, with values further from zero indicating greater distinction between classes. This distribution inherently captures prediction uncertainty, encompassing both analytical and spectral variations. The NCD offers several advantages, including, (i) maps high-dimensional spectra to an interpretable scalar value, (ii) captures non-linear relationships between spectral features, (iii) provides an unbounded measure that effectively captures subtle variations in spectral quality., (iv) learns relevant patterns from training data.

To ensure robust model selection and unbiased performance estimation, nested cross-validation (NCV) is recommended [14,15]. This involves splitting the dataset into a calibration set for training and finetuning, and an external validation set for independent assessment. The calibration set undergoes k-fold cross-validation, yielding k different models that collectively assess performance stability. This approach prevents overfitting and yields reliable estimates of the model’s generalization ability.

## 4 Evaluation of distances using MALDI-TOF spectra

### 4.1 Spectral dataset and scenarios for spectral variability

To demonstrate the above described concept and evaluate the distances, we make use of Matrix-Assisted Laser Desorption/Ionization Time-of-Flight (MALDI-TOF) mass spectrometry spectra for Methicillin-resistant *Staphylococcus aureus* (MRSA) and Methicillin-susceptible *Staphylococcus aureus* (MSSA) and formulate three cases as illustrated in Figure 1.

**Figure 1.**
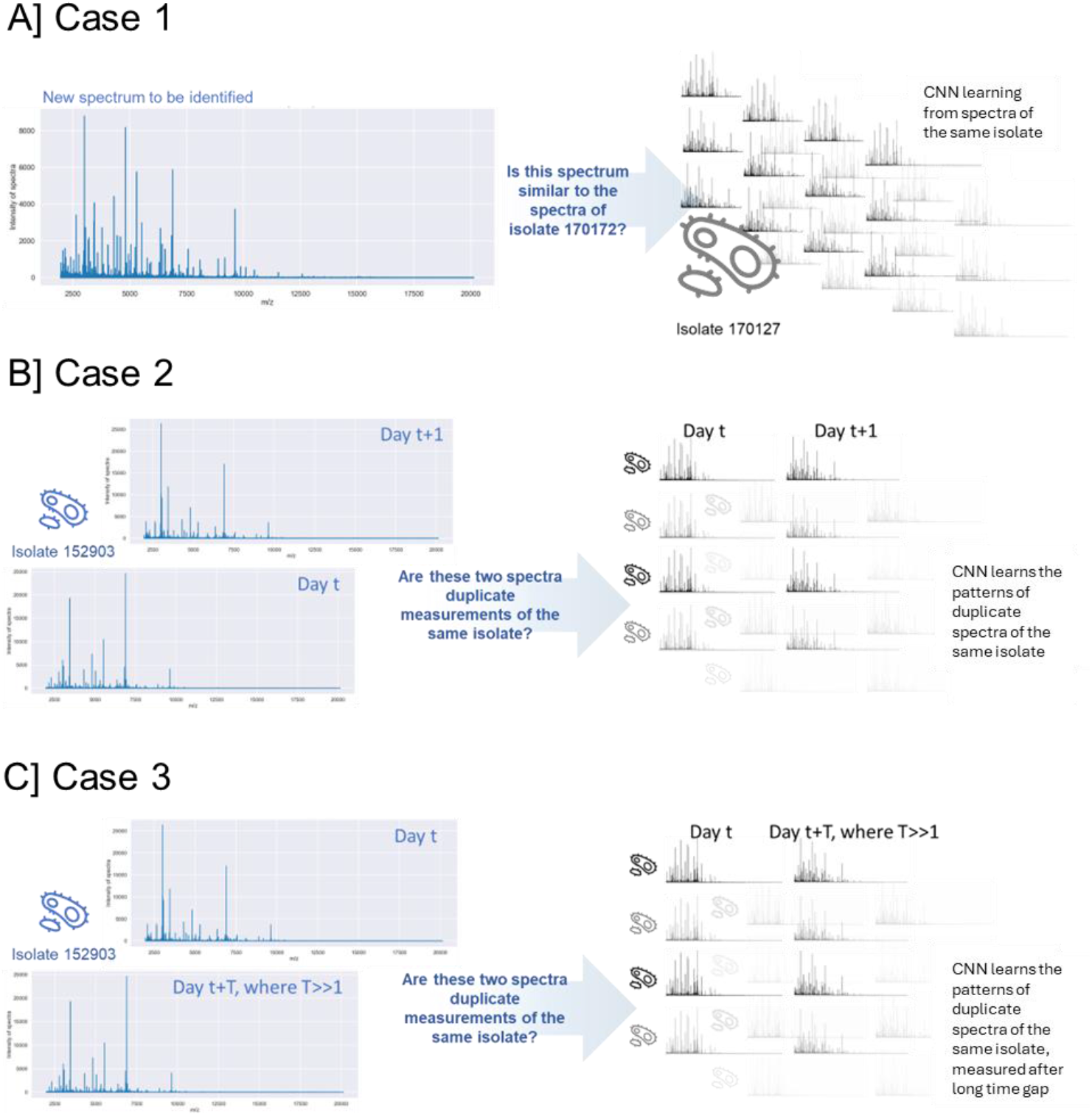
Illustration of 3 cases for the data experiments performed. (A) Case 1 for within-run variability. (B) Case 2 for day-to-day comparisons, (C) Case 3 for cross-run (long time) robustness.

The spectra underwent baseline correction removed background noise by estimating (rolling median) and subtracting a baseline from each spectrum. The available dataset of 923 MALDI-TOF mass spectrometry measurements, 658 MSSA and 265 MRSA, were used. This dataset fortuitously provided an opportunity to exploit three distinct analytical scenarios. We systematically constructed three test cases that would build an understanding of spectral relationships (see Figure 1). By strategically partitioning the available measurements, we created complementary scenarios progressing from fundamental within-run variability (Case 1) through temporal comparisons (Case 2) to cross-run (long time) robustness (Case 3). This fortuitous arrangement allowed us to examine spectral discrimination at different levels of experimental variability. The data compilation was done by looking at the isolate numbers, type of isolate and whether it was a repeat measurement. Note that this study does not compare MRSA versus MSSA characteristics; rather, each case examines within-type spectral variability across different temporal and experimental conditions.

Case 1 investigates whether spectral patterns can distinguish repeat measurements of a single isolate from different isolates. The dataset comprises 92 same-day spectra from one MSSA isolate and 92 spectra from 41 different MSSA isolates. This allows evaluation of same-day measurement detection capability.

Case 2 explores more subtle spectral differences, specifically whether spectra from the same isolate measured one day apart remain distinguishable from different isolates measured simultaneously. The dataset contains 40 MRSA isolates, each measured twice one-day apart.

Case 3 examines long-term spectral variability using 45 MSSA isolates measured twice, approximately three months apart. The analysis determines whether same-isolate spectra maintain closer relationships than different-isolate spectra measured on the same day, despite the extended time gap.

For each case, we evaluated the spectral relationships using NCD and MD. NCD was derived by training convolutional neural network (CNN) models to distinguish between within-isolate and between-isolate spectral differences, offering a machine learning perspective on spectral discrimination. This dual approach strengthened our assessment of spectral relationships across increasing levels of experimental variability.

### 4.2 Results for MD for different scenarios of spectral variability

Figure 2 shows the results for MD evaluated for the three cases. In Case 1, using a dataset of 92 spectra from one MSSA isolate (class 1) and 41 different MSSA isolates (class 2), we evaluated MD using covariance matrices derived from both same-isolate measurements (Σ_1_) and different-isolate measurements (Σ_2_). For 92+41 = 133, 8778 different pairs of distances were evaluated. Using 60 spectra for calculating Σ_1_, the expected MD between corresponding spectra was empirically set to 12, rather than using the theoretical value of 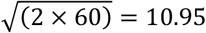. Since raw distance values alone were insufficient for isolate classification, we established a decision threshold to categorize samples into class 1 (same isolate) or class 2 (different isolate). The threshold of 13.77 (p = 0.95) was determined using an empirical expected MD value of 12 multiplied with the inverse chi-squared distribution. This was calculated as 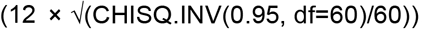, where CHISQ.INV(0.95, 60) = 79.08.

**Figure 2.**
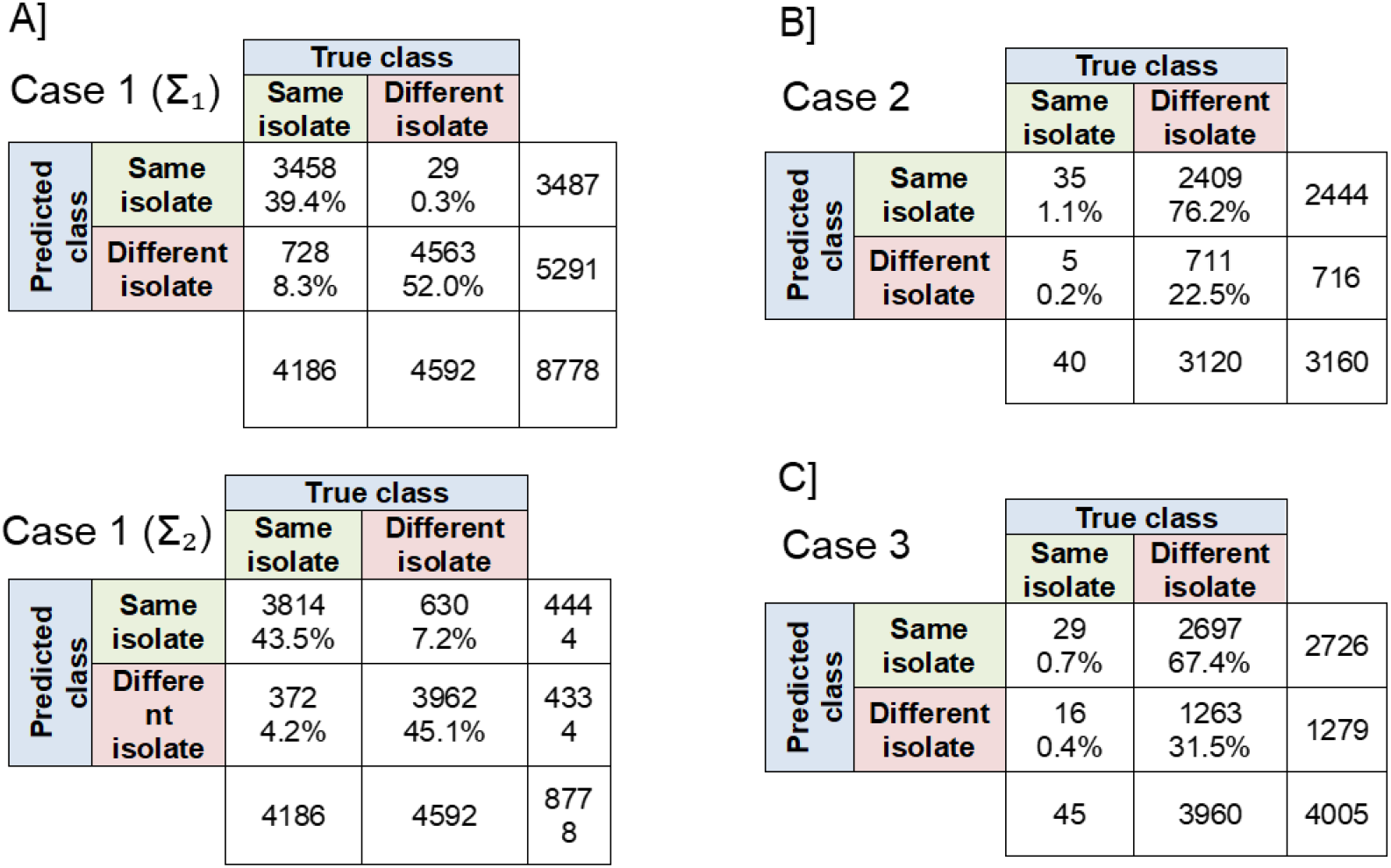
Confusion matrices for the three cases for calculation of Mahalanobis distances: (A) case 1, (B) case 2 and (C) case 3.

Out of these 8778, a total of 4563 pairs of spectra are classified correctly as the different isolate and a total of 3458 pairs of spectra are classified correctly as same isolates.

Similarly, 41 spectra of class 2 were split into groups of 30 and 11 spectra and the former is used for the calculation of Σ_2_. In this case, an empirical decision threshold of 7 was chosen, resulting in 3814 pairs of spectra correctly identified as same isolate and 3962 pairs of spectra correctly identified as different isolate. Together, these results show that classification based on the MD based metrics is possible, but with some false positives (8.3% and 4.2%) and false negatives (0.3% and 7.2%).

Moving to Case 2 using 40 pairs of MRSA spectra on a “day t” were split into groups of 30 and 10 spectra, respectively. The former is used for the calculation of Σ. Remaining 10 spectra from class 2, are set aside and not used in the calculation of Σ. Additional 40 spectra of the same isolates measured on day t+1 from the same isolates were used in the pairwise distance calculations.

MD analysis using the day-t covariance matrix achieved very few correct classifications (35+711=746 out of 3160) with a 9.35 threshold. The large increase in false positive rate (76.2%) compared to the previous scenario suggests limitations in MD’s ability to capture complex spectral relationships.

Lastly, in Case 3, using 45 pairs of MSSA spectra with maximum expected variation, MD correct classifications remained poor (29+1263=1292 out of 4005), with a 67.4% false positive rate. This significant decrease in performance indicates MD’s limitations in handling highly variable spectral relationships.

When the number of variables approaches or exceeds the number of observations, the sample covariance matrix becomes ill-conditioned or singular. This fundamentally compromises the calculation of MD, as the covariance matrix inverse becomes unstable or undefined. The issue is particularly acute in spectroscopic data where the number of wavelengths (variables) typically far exceeds the number of spectra (observations). This ‘curse of dimensionality’ necessitates either dimensionality reduction, regularization of the covariance matrix, or alternative distances that remain stable in high-dimensional spaces. NCD offers an appealing alternative, circumventing these dimensionality constraints through learned feature representations.

### 4.3 Results for NCD for different scenarios of spectral variability

In Case 1, a CNN was trained with 5-fold NCV for classifying spectra corresponding to class 1 (same isolate) and class 2 (different isolates). The external validation set comprised of 22 spectra from class 1 and 11 spectra from class 2. The NCD demonstrated superior discrimination capability, achieving 100% classification accuracy in both internal (n=70) and external (n=33) validation set (considering zero as decision threshold) (see Figure 3A). The clear separation in NCD indicates robust feature extraction beyond linear distance. Absence of misclassifications establishes NCD as a reliable distance value for discriminating spectra under repeatability conditions, providing a good basis for more complex temporal comparisons.

**Figure 3.**
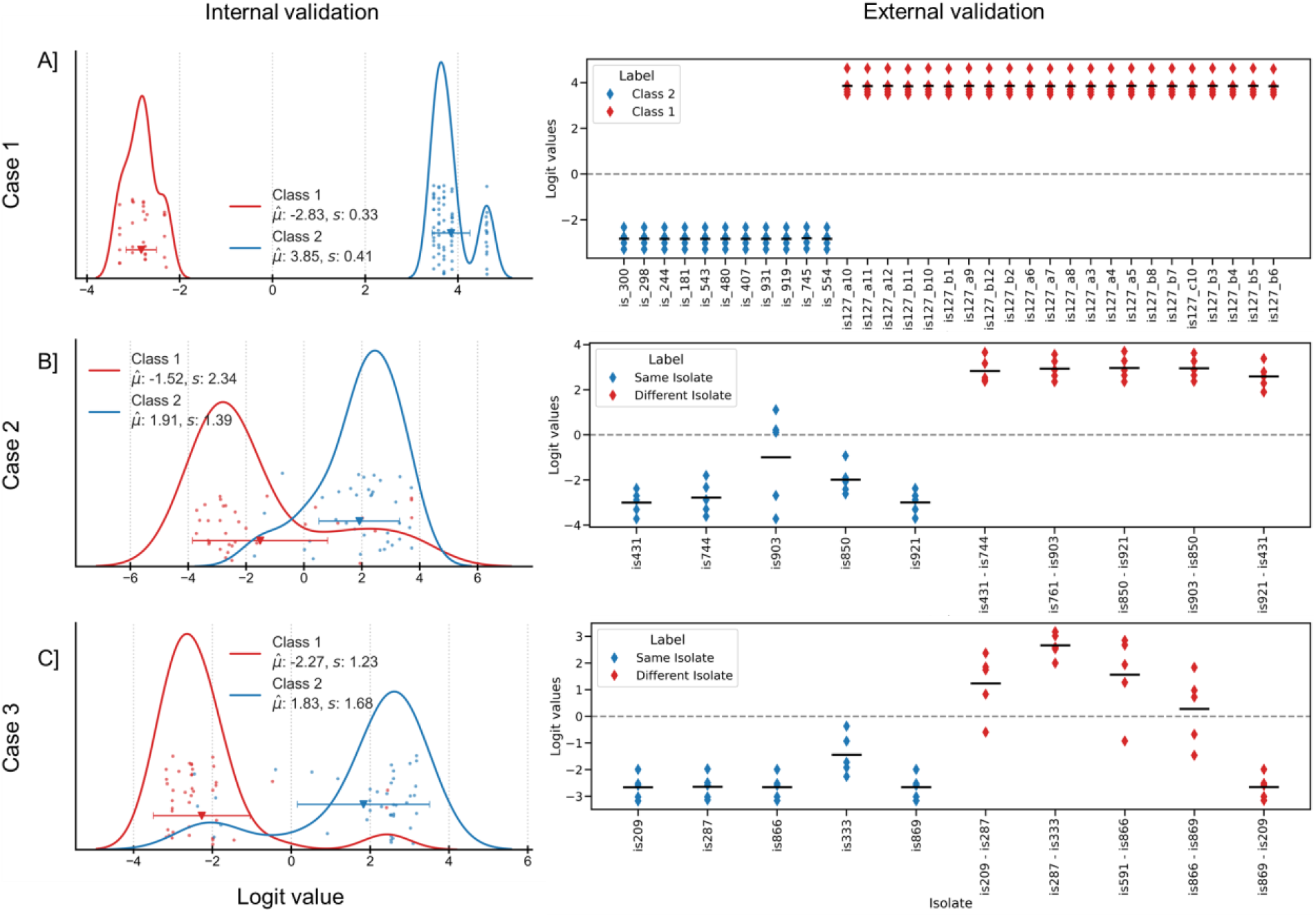
Summary of NCD results for the three cases: A) case 1, B) case 2 and C) case 3. Left panels show distribution of logit values for internal validation. The right panels show the distribution for external validation samples.

Turning to Case 2, here the differences in the spectra were used to construct a set of data for the training a CNN. On one hand, the difference between the replicates (measured a day apart) were used to generate a collection of forty samples that each represented data from the same isolate. On the other hand, for the “counter class”, spectra from randomly selected isolates were subtracted, resulting in the creation of another dataset containing forty samples. For the purposes of training and verifying the model, only 70 of the initial 80 samples were utilized. They were divided into five equal folds, each containing a total of 14 samples. The remaining 10 samples were utilized for the purpose of conducting an external validation. Figure 3B shows the logit values for the external validation dataset. All spectra of the classes 1 (difference between replicate of the same isolate) and 2 (difference between two different isolate) were classified correctly. The CNN models, trained on spectral differences, maintained stronger performance with 81% of the spectra correctly classified in internal validation (n=70) and 100% in external validation (n=10). One isolate “is903” with its large distribution of the logit values, close to the decision threshold, appears to be a “challenging sample”.

Lastly, for Case 3, another CNN was trained similar to the one applied for the previous case where class 1 comprised 45 samples (differences between spectra of the same isolate measured in two different rounds) and class 2 comprised 45 samples (differences between spectra from two isolates measured during the same round). 10 samples were separated for external validation and the remaining 80 samples were organized in 5 folds of 16 samples each. Figure 3C shows the distribution of NCD, with 90% accuracy in internal validation (n=80) and 80% in external validation (n=10). The logit value distribution reveals-maintained class separation, though with reduced margin compared to simpler scenarios, indicating systematic handling of complex spectral relationships.

Overall, these results demonstrate that while both distances (MD and NCD) can effectively discriminate spectral differences under controlled conditions, NCD maintains more robust performance as spectral complexity increases. The progressive decline in MD performance contrasted with the relatively stable CNN accuracy suggests fundamental differences in how these methods handle spectral variation.

## 5 Discussion

The implementation of neural networks for spectral data quality assessment represents a significant advancement in NTMs. By enabling automated detection of anomalous patterns, this approach establishes objective criteria for spectral database curation, addressing a crucial gap in quality assurance protocols. Our proposed framework using decision scores addresses the question of how the error can be characterized.

The quantitative logit values obtained as the classification output has not only the role for performance evaluation but also for data quality evaluation as shown here [7]. They serve as a distance to numerically and graphically interpret the similarity or dissimilarity between the spectra. The NCD approach presented here provides a foundation for developing robust quality metrics for analytical techniques using spectral data.

The NCD approach demonstrates several key advantages over traditional methods. Most notably, while the Mahalanobis distance assumes a static covariance structure (Σ), NCD adapts to temporal changes through its learned embedding function *ϕ*. This adaptability is evidenced by superior classifications in the example application, where NCD’s capacity to capture subtle spectral variations significantly outperformed the MD approach.

With multivariate data, a classical match factor approach can be adopted which reduces this complexity to a single similarity value. This approach however presents several limitations including but not limited to (a) the reduction to a single value obscures the multidimensional nature of spectral variations, and (b) fails to account for measurement conditions (repeatability r and reproducibility R), sampling or sample preparation variations. The NCD approach tries to overcome these limitations. The transition from single-value similarity metrics to NCDs allows decomposition into hierarchical sources of variability represents a significant advancement in NTM quality assurance. Quality assurance tools will benefit greatly from such metrics for quantifying discrepancies between spectra.

The integration of NCD with traditional control charts demonstrates particular promise, offering a robust framework for continuous quality monitoring that extends beyond conventional targeted approaches. The application of control charts to non-targeted analysis introduces a systematic approach to quality control that was previously challenging to implement. By monitoring NCD over time, laboratories can now detect subtle deviations in spectral patterns that might indicate instrumental drift or sample preparation inconsistencies. This capability is particularly valuable given the complex nature of NTMs, where traditional quality control parameters may be insufficient or inappropriate.

The proposed approach can be used for outlier identification as well. By leveraging the distances between spectra, the method provides a quantitative basis for identifying anomalous measurements. This objective approach to outlier detection strengthens the reliability of spectral databases and supports more confident data interpretation in NTMs.

The comparison of spectral preprocessing techniques using NCD represents another significant use case. This approach provides an objective means of evaluating preprocessing strategies, enabling laboratories to optimize or standardize their analytical workflows based on quantitative evidence rather than empirical observation.

Perhaps most significantly, this work establishes a foundation for new proficiency testing schemes specifically designed for NTM. The use of NCD offers a novel approach to inter-laboratory comparisons, potentially enabling standardized quality assessment across different laboratories and instruments. This development addresses a long-standing challenge in the field of non-targeted analysis, where traditional proficiency testing approaches have proven inadequate.

These findings extend beyond MALDI-TOF MS to the broader field of non-targeted methods, where similar challenges in spectral comparison and quality assurance persist. The NCD approach provides a universal framework for developing robust quality metrics across analytical techniques using spectral data. These advancements collectively suggest a path toward more robust and reliable NTMs, with implications for fields ranging from environmental monitoring to food safety analysis. Future work should focus on the standardization of these approaches and their integration into routine laboratory practice.

## 6 Conclusion

This study demonstrates a significant advancement in spectral variation methodology for NTM. While traditional approaches like MD provide acceptable performance under controlled conditions, their effectiveness diminishes with increasing spectral complexity. The NCD approach exhibits superior classification accuracy across varying analytical challenges. And hence, it serves multiple purposes: it enables quantitative assessment of spectral similarities, provides a foundation for quality metrics, and offers insights into method limitations. Our findings establish that modern machine learning approaches can effectively complement classical statistical methods in analytical quality assurance.

Of particular significance is the framework’s potential for practical implementation in analytical laboratories. The methodology developed here can be readily adapted for routine quality control, method validation, and stability assessment in non-targeted analysis. Future applications could extend beyond mass spectrometry to other analytical techniques where spectral comparison plays a crucial role. This work represents a step toward standardizing quality assessment in NTM, though further validation across different analytical platforms and applications remains an important area for future research.

